# COVID-19: Viral-host interactome analyzed by network based-approach model to study pathogenesis of SARS-CoV-2 infection

**DOI:** 10.1101/2020.05.07.082487

**Authors:** Francesco Messina, Emanuela Giombini, Chiara Agrati, Francesco Vairo, Tommaso Ascoli Bartoli, Samir Al Moghazi, Mauro Piacentini, Franco Locatelli, Gary Kobinger, Markus Maeurer, Alimuddin Zumla, Maria R. Capobianchi, Francesco Nicola Lauria, Giuseppe Ippolito, COVID 19 INMI Network Medicine for IDs Study Group

## Abstract

**Background:** Epidemiological, virological and pathogenetic characteristics of SARS-CoV-2 infection are under evaluation. A better understanding of the pathophysiology associated with COVID-19 is crucial to improve treatment modalities and to develop effective prevention strategies. Transcriptomic and proteomic data on the host response against SARS-CoV-2 still have anecdotic character; currently available data from other coronavirus infections are therefore a key source of information.

**Methods:** We investigated selected molecular aspects of three human coronavirus (HCoV) infections, namely SARS-CoV, MERS-CoV and HCoV-229E, through a network based-approach. A functional analysis of HCoV-host interactome was carried out in order to provide a theoretic host-pathogen interaction model for HCoV infections and in order to translate the results in prediction for SARS-CoV-2 pathogenesis.

The 3D model of S-glycoprotein of SARS-CoV-2 was compared to the structure of the corresponding SARS-CoV, HCoV-229E and MERS-CoV S-glycoprotein. SARS-CoV, MERS-CoV, HCoV-229E and the host interactome were inferred through published protein-protein interactions (PPI) as well as gene co-expression, triggered by HCoV S-glycoprotein in host cells.

**Results:** Although the amino acid sequences of the S-glycoprotein were found to be different between the various HCoV, the structures showed high similarity, but the best 3D structural overlap shared by SARS-CoV and SARS-CoV-2, consistent with the shared ACE2 predicted receptor. The host interactome, linked to the S-glycoprotein of SARS-CoV and MERS-CoV, mainly highlighted innate immunity pathway components, such as Toll Like receptors, cytokines and chemokines.

**Conclusions:** In this paper, we developed a network-based model with the aim to define molecular aspects of pathogenic phenotypes in HCoV infections. The resulting pattern may facilitate the process of structure-guided pharmaceutical and diagnostic research with the prospect to identify potential new biological targets.

## Background

In December 2019, a novel coronavirus (SARS-CoV-2) was first identified as a zoonotic pathogen of humans in Wuhan, China, causing a respiratory infection with associated bilateral interstitial pneumonia. The disease caused by SARS-CoV-2 was named by the World Health Organization as COVID-19 and it has been classified as a global pandemic since it has spread rapidly to all continents. As of March 30, 2020, there have been 634 835 confirmed COVID-19 cases worldwide with 29 957 deaths reported to the WHO [1]. Whilst clinical and epidemiological data on COVID-19 have become readily available, information on the pathogenesis of the SARS-CoV-2 infection has not been forthcoming [2].The transcriptomic and proteomic data on host response against SARS-CoV-2 is scanty and not effective therapeutics and vaccines for COVID-19 are available yet.

Coronaviruses (CoVs) typically affect the respiratory tract of mammals, including humans, and lead to mild to severe respiratory tract infections [3]. Many emerging HCoV infections have spilled-over from animal reservoirs, such as HCoV-OC43 and HCoV-229E which cause mild diseases such as common colds [4, 5]. During the past 2 decades, two highly pathogenic HCoVs, severe acute respiratory syndrome coronavirus (SARS-CoV) and Middle East respiratory syndrome coronavirus (MERS-CoV), have led to global epidemics with high morbidity and mortality [6]. In this period, a large amount of experimental data associated with the two infections has allowed to better understand molecular mechanism(s) of coronavirus infection, and enhance pathways for developing new drugs, diagnostics and vaccines and identification of host factors stimulating (proviral factors) or restricting (antiviral factors) infection remains poorly understood [7]. Structures of many proteins of SARS-CoV and MERS-CoV, and biological interactions with other viral and host proteins have been widely explored; through experimental testing of small molecule inhibitors with anti-viral effects [8, 9]. ACE2, expressed in type 2 alveolar cells in the lung, has been identified as receptor of SARS-CoV and SARS-CoV-2, while dipeptidyl peptidase DPP4 was identified as the specific receptor for MERS-CoV [10, 11].

The investigation of structural genomics and interactomics of SARS-CoV-2 can be implemented through an integrated bioinformatics approach. Structural analysis of specific SARS-CoV-2 proteins, in particular Spike glycoproteins (S-glycoproteins), and their interactions with human proteins, can guide the identification of the putative functional sites and help to better define the pathologic phenotype of the infection. This functional interaction analysis between the host and other HCoVs, combined with an evolutionary sequence analysis of SARS-CoV-2, can be used to guide new treatment and prevention interventions.

We investigated here biologically and clinically relevant molecular targets of three human coronaviruses (HCoV) infections using a network based approach. A functional analysis of HCoV-host interactome was carried out in order to provide a theoretic host-pathogen interaction model for HCoV infections, and to predict viable models for SARS-CoV-2 pathogenesis. Three HCoV causing respiratory diseases were used as the model targets, namely: SARS-CoV, that shares with SARS-CoV-2 a strong genetic similarity, including MERS-CoV, and HCoV-229E.

## Methods

### Comparative reconstruction of S-glycoprotein in HCoVs

The reconstruction of virus-host interactome was carried out using the RWR algorithm to explore the human PPI network and the multilayer PPI platform enriched with gene expression data sets. 259 sequences of CoVs, infecting different animal hosts (Table S1 in Additional file 1), were downloaded by GSAID and NCBI database in order to evaluate the variability in the S gene. SARS-CoV, HCoV-229E and MERS-CoV and other CoV full genome sequence groups were aligned with MAFFT [12], synonymous and non-synonymous mutations, and amino acid similarity were calculated using the SSE program with a sliding windows of 250 nucleotides and a pass of 25 nu [13]. A homology model was built for the amino acid sequences of the S-glycoprotein, derived from the full genome sequence obtained at “SARS-CoV-2/INMI1/human/2020/ITA” (MT066156.1). The Swiss pdb server was used to construct three-dimensional models for the S-glycoprotein of SARS-CoV-2 [14]. Among proteins with a 3D structure, the best match with the “SARS-CoV-2/INMI1/human/2020/ITA” sequence was the 6VSB.1, that was evaluated considering the identity of two amino acid sequences and the QMEN value included in Swiss pdb server. The model of a single chain was overlapped with the three-dimensional structure of S-glycoprotein single chain belonging to SARS-CoV (5WRG), HCoV-229E (6U7H.1) and MERS-CoV (5×59), using Chimera 1.14 [15]. In order to better evaluate the conservation of the sequence in each site, all sequences were aligned with MAFFT and the topology of all structures were compared. The detailed description of the reconstruction of S-glycoprotein structure is reported in the Appendix.

### PPI and Gene Co-expression Network

Network analysis, based on protein-protein interactions and gene expression data, was performed in order to view all possible virus-host protein interactions during the HCoV infections. Since the SARS-CoV-2 genome exhibits substantial similarity to the SARS-CoV genome [16] and subsequently also the proteome [17], we hypothesized that several molecular interactions that were observed in the SARS-CoV interactome will be preserved in the SARS-CoV-2 interactome. Virus-Host interactomes (SARS-CoV, MERS-CoV, HCoV-229E) were inferred through published PPI data, using two publicly accessible databases (STRING Viruses and VirHostNet), as well as published scientific reports with a focus on virus-host interactions [18–20]. As a next step, the virus-host PPI list, extracted in this first step, was merged with additional PPI databases, i.e. BioGrid, InnateDB-All, IMEx, IntAct, MatrixDB, MBInfo, MINT, Reactome, Reactome-FIs, UniProt, VirHostNet, BioData, CCSB Interactome Database, using R packages PSICQUIC and biomaRt [21, 22]. In total, a large PPI interaction database was assembled, including 13,020 nodes and 71,496 interactions.

The gene expression data set was built from the Protein Atlas database, using tissue and cell line data [23]. To identify the most likely interactions, and to obtain functional information, Random walk with restart (RWR), a state-of-the-art guilt-by-association approach by R package RandomWalkRestartMH [24] was used. It allows to establish a proximity network from a given protein (seed), to study its functions, based on the premise that nodes related to similar functions tend to lie close to each other in the network. For each node, a score was computed as measure of proximity to the seed protein. S-glycoproteins of SARS-CoV, MERS-CoV and HCoV-229E were used as seed in the application of the RWR algorithm.

### Functional Enrichment Analysis

To evaluate functional pathways of proteins involved in host response, gene enrichment analysis was performed, using Kyoto Encyclopedia of Genes and Genomes (KEGG) human pathways and Gene Ontology databases. Network representation from the gene enrichment analysis was performed using ShinyGO v0.61[25]. The statistical significance was obtained, calculating the False Discovery Rate (FDR).

## Results

### Structure of S-glycoprotein CoVs

To evaluate the diversity along the full genome, pairwise distance was calculated on 259 HCoV genomes. Diversity was distributed along the entire CoV genome, with the most conserved region located in Orf1ab, as expected, while the spike gene region exhibited a rather high diversity (Figure S1 in Appendix).

Consequently, the analysis was focused on the S-glycoprotein, as a key virus component involved in host interaction [26]. A 3D model of S-glycoprotein of the SARS-CoV-2 sequence (MT066156.1) was built on the sequence obtained at Laboratory of Virology, National Institute of Infectious Diseases “L. Spallanzani” IRCCS, using Swiss pdb viewer server (Figure S2 a/b in Appendix). The SARS-CoV-2 S-glycoprotein structure was then compared to other HCoVs as shown in Figure S2 in Appendix. The S-glycoprotein structures of the various HCoV were very similar overall. In particular, a strong similarity was shown in the RBD (nCov: residues 319−591) [27], and this was most evident for the comparison between SARS-CoV-2 and SARS-CoV, which share the same cell receptor (ACE2). The amino acid differences among the S-glycoproteins of the selected HCoVs (SARS-CoV-2, MERS-CoV, SARS-CoV, HCoV-229E) are shown in Figure S3 in Appendix, where a lower topology similarity was observed with HCoV-229E S-glycoprotein, which binds a different host receptor.

Overall, the pattern arising from such comparison was consistent with specific host receptors, as well as with different host reservoirs and ancestry [28].

### Human CoV and host interactome

An interactome map was built to highlight biological connections among S-glycoprotein and the human proteome. Using the analysis pipeline described in the methods, a large PPI interaction database was assembled, including 13,020 nodes and 71,496 interactions between human host and the three selected viruses (SARS-CoV, MERS-CoV and HCoV-229E).

The interactome reconstruction was obtained with the RWR analysis, finding 200 closest proteins to seed, or S-glycoproteins of HCoV-229E, SARS-CoV and MERS-CoV (Figures S4-S6 in Appendix). In Table S2, S3 and S4 in Additional file 1, lists of genes selected by RWR algorithm for HCoV-229E, SARS-CoV and MERS-CoV, along with proximity score were reported. In order to further dissect the S-glycoprotein-host interactions, enrichment analysis was carried out with Reactome and KEGG databases. Reactome pathway enrichment analysis revealed biological pathways of DNA repair, transcription and gene regulation for the HCoV-229E S-glycoprotein, with high significance (FDR <0.01%). KEGG pathway enrichment analysis revealed ubiquitin mediated proteolysis as the most significant pathway (FDR <0.01%), as well as altered cellular proliferation pathways, associated with other viral infections (Hepatitis B, measles, Epstein-Barr virus infection and Human T-cell leukemia virus 1 infection) as well as with carcinogenesis (Figure 1). Next, the RWR algorithm was applied to a multilayer network built on the PPI interactome and on the Gene Coexpression (COEX) network, again with S-glycoprotein of HCoV-229E as seed. The results highlighted a set of genes that are connected in both PPI and COEX analysis, including ANPEP, RAD18, APEX, POLH, APEX1, TERF2, RAD51, CDC7, USP7, XRCC5, RAD18, FEN1, PCNA, all associated to the GO biological process category of DNA repair (FDR< 0.0001%) (Figure 2). The same analyses were conducted for SARS-CoV and MERS-CoV.

**Figure 1.**
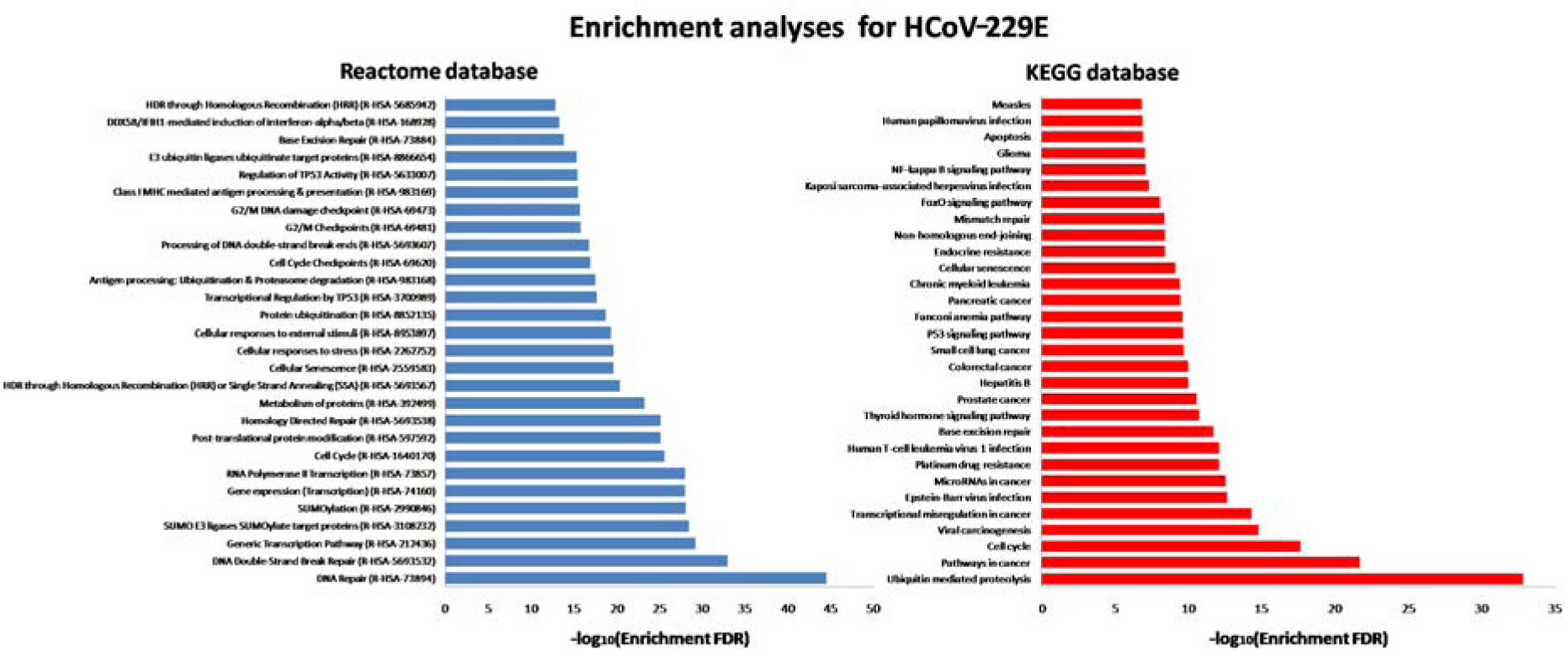
KEGG human pathway and Reactome pathways enrichment analysis for 200 proteins identified by RWR algorithm using S-glycoprotein of HCoV-229E.

**Figure 2.**
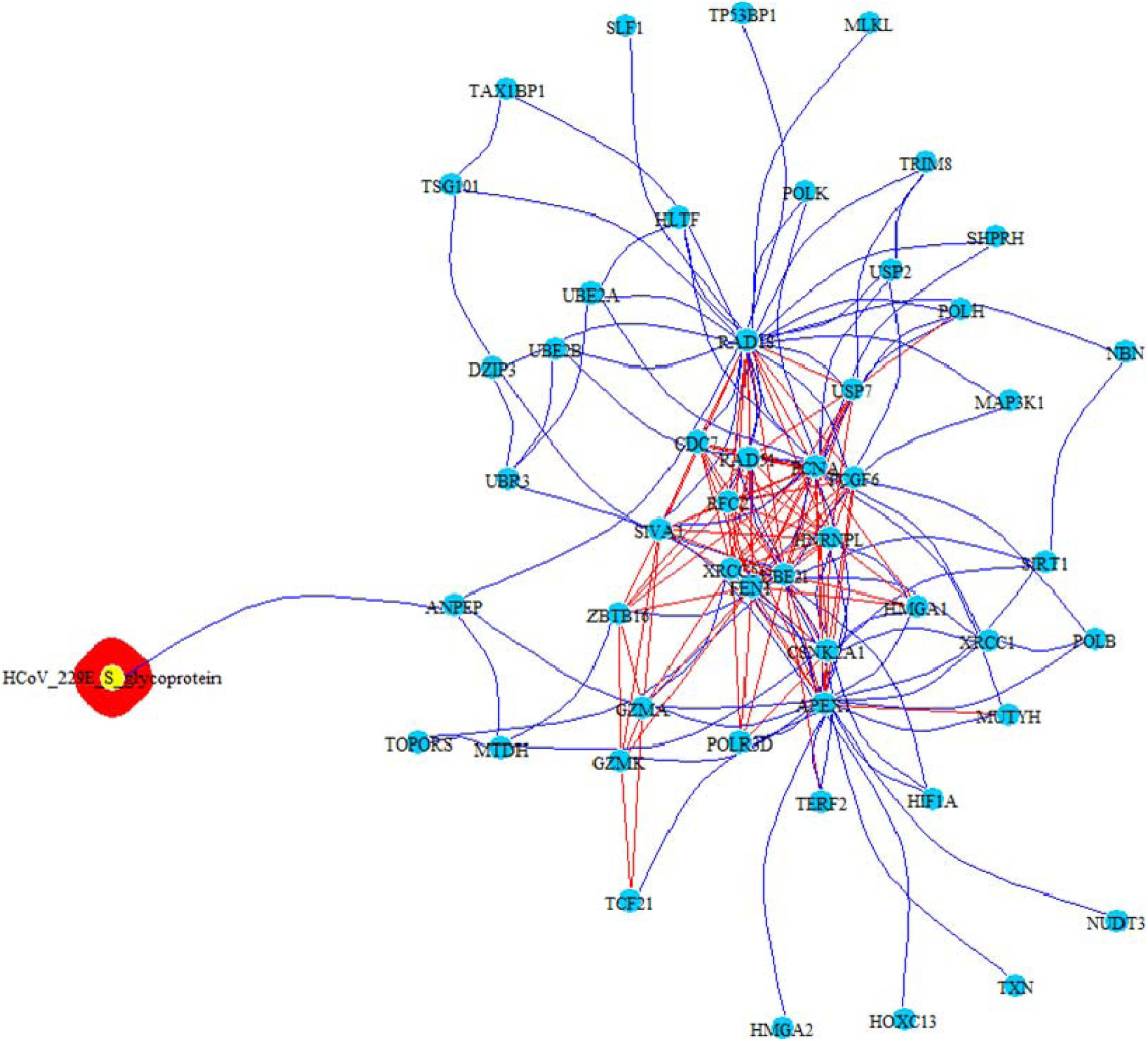
PPI-COEX multilayer analysis, based on human PPI interactome and Gene Coexpression network, with top 50 closest proteins/genes identified by RWR, using S-glycoprotein of HCoV-229E. Edges in blue represent PP interactions, while red edges are coexpressions.

The Reactome pathway enrichment analysis for the SARS-CoV revealed S-glycoprotein connection with early activation of innate immune system, such as the Toll Like Receptor Cascade and TGF-β, with a strong significance (FDR < 0.0001%), while the KEGG pathway enrichment analysis revealed an association with cellular proliferation, TGF-β and other infection-related pathways (FDR<0.0001%) (Figure 3). The PPI-COEX multilayer analysis highlighted a set of genes that are connected in both PPI and COEX analysis, i.e.CLEC4G, CLEC4M, CD209, ACE2, RPSA, all associated to the GO biological process category of SARS-CoV entry into host cell (FDR < 0.01%) (Figure 4).

**Figure 3.**
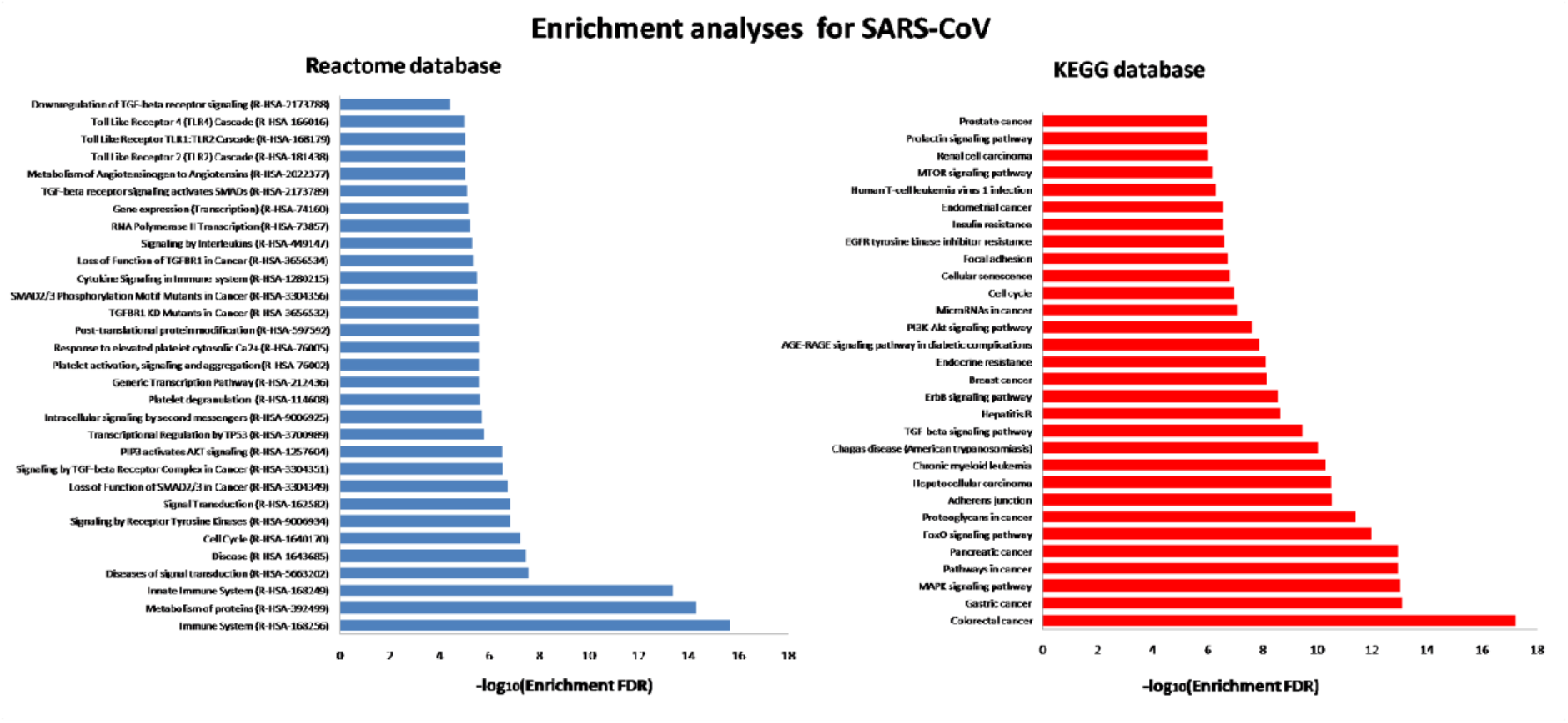
KEGG human pathway and Reactome pathways enrichment analyses for 200 proteins identified by RWR algorithm using S-glycoprotein of SARS-CoV.

**Figure 4.**
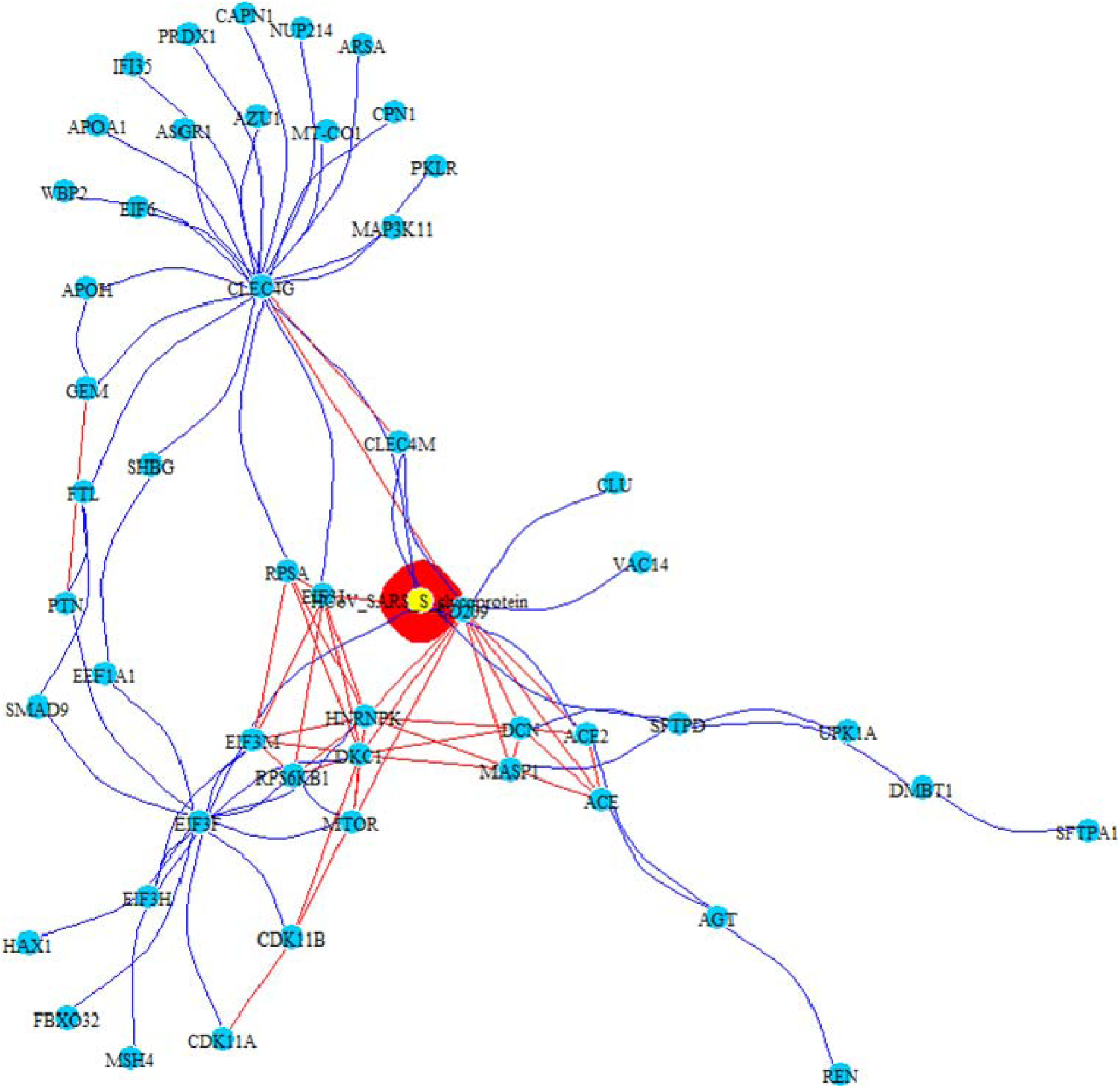
PPI-COEX multilayer analysis based on human PPI interactome and Gene Coexpression network, with top 50 closest proteins/genes identified by RWR, using S-glycoprotein of SARS-CoV. Edges in blue represent PP interactions, while red edges are coexpressions.

In MERS-CoV, the Reactome pathway enrichment analysis showed a strong association with membrane signals activated by GPCR ligand binding (FDR < 0.0001%), and chemokine/chemokine receptor pathways. Consistent results were obtained with KEGG pathway enrichment, that highlighted cytokine-cytokine receptor and chemokine signalling pathways (FDR<0.0001%) (Figure 5). Finally, PPI-COEX multilayer analysis evidenced, for both PPI and COEX, CCR4, CXCL2, CXCL10, CXCL9, PF4, PF4V1, CCL11, CXCL11, XCL1, CXCR4 and CXCL14, all genes identified by the GO biological processes involved in the chemokine cascade (FDR < 0.0001%), in line with the results obtained with enrichment analyses (Figure 6).

**Figure 5.**
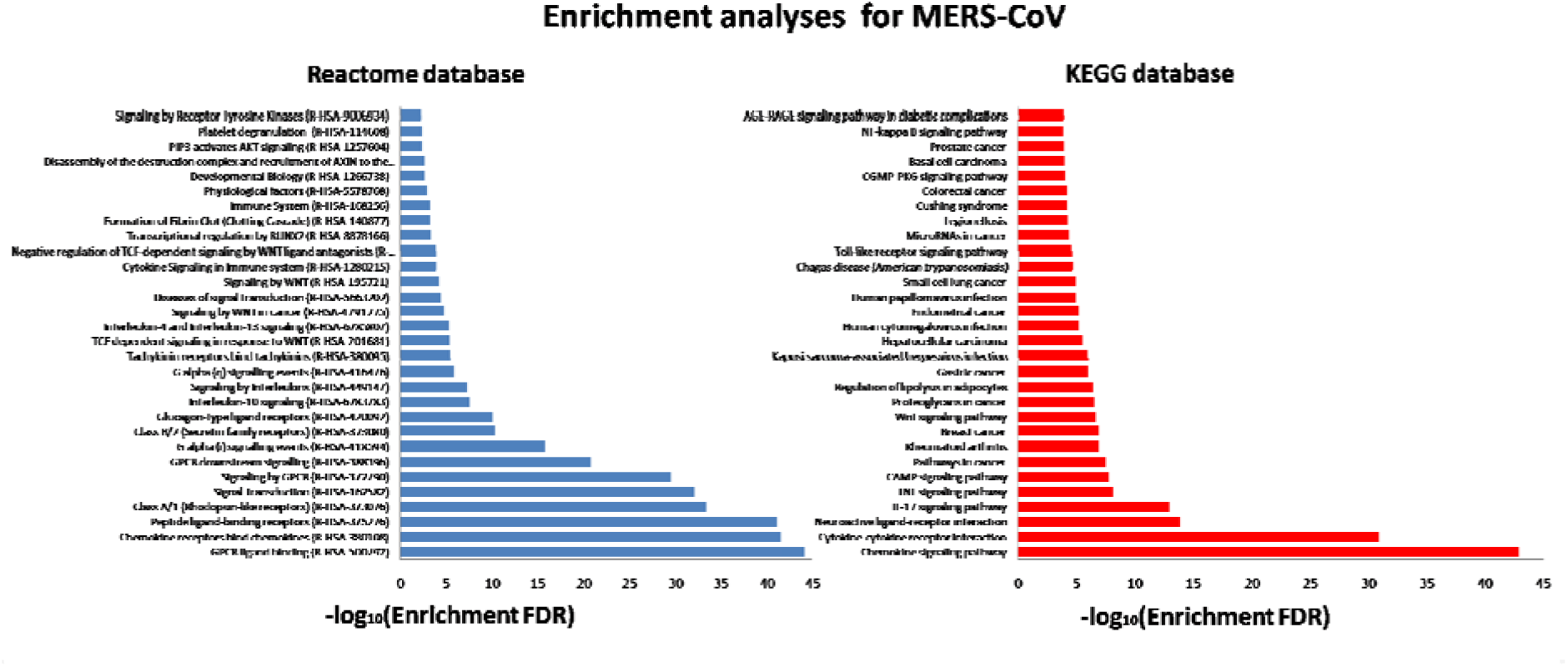
KEGG human pathway and Reactome pathways enrichment for 200 proteins identified by RWR algorithm using S-Glycoprotein of MERS-CoV.

**Figure 6.**
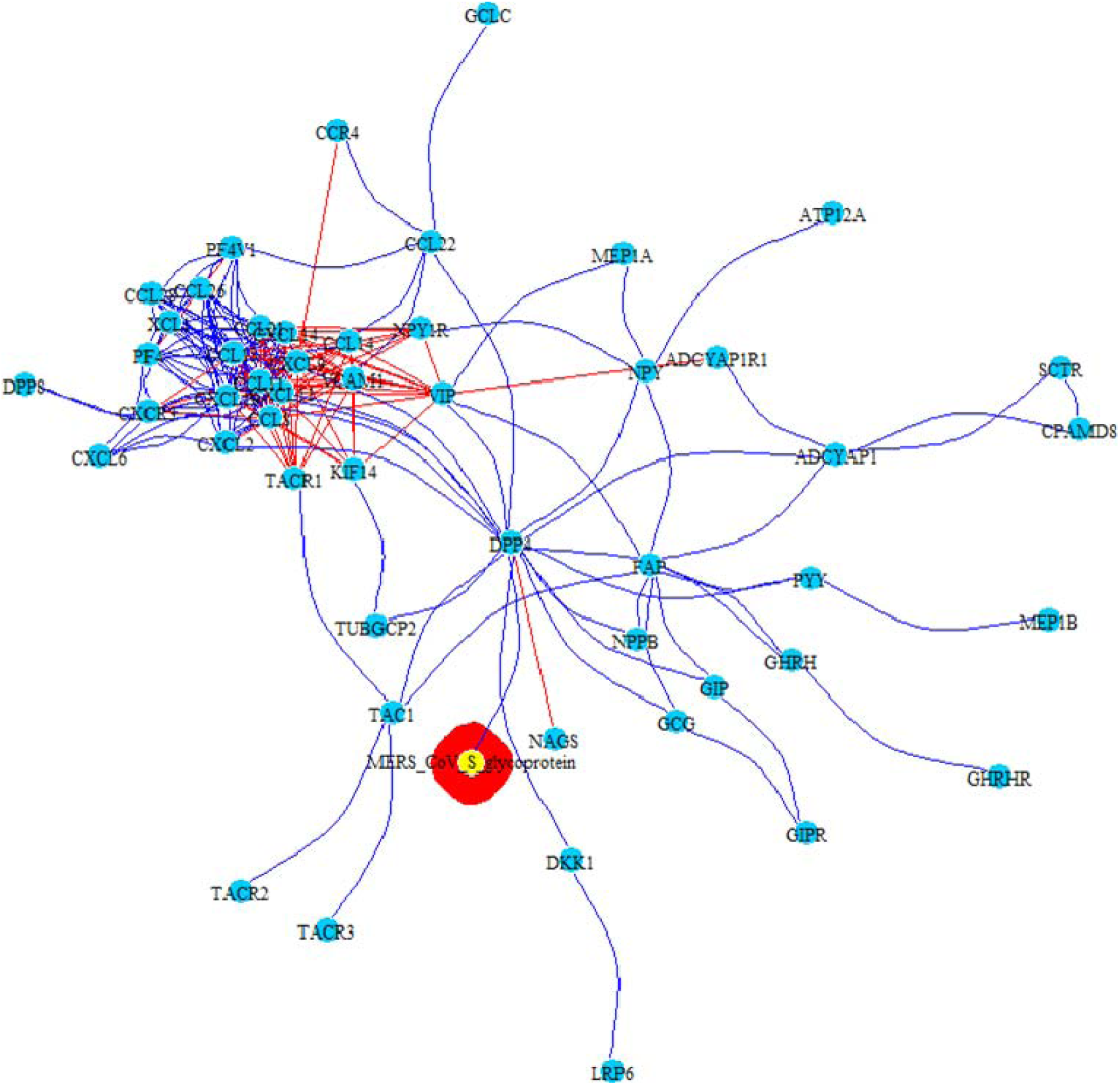
PPI-COEX mulilayer analysis based on human PPI interactome and Gene Coexpression network, with top 50 closest proteins/genes identified by RWR, using S-glycoprotein of MERS-CoV. Edges in blue represent PP interactions, while red edges indicate coexpressions.

## Discussion

### In-depth comparative analysis of S-glycoprotein

We applied network analysis, based on protein-protein interactions and gene expression data, in order to describe the interactome of the coronavirus S-glycoprotein and host proteins, with the aim to better understand SARS-CoV-2 pathogenesis. A preliminary structural analysis was conducted on the S-glycoprotein of SARS-CoV-2 as compared to the other 3 HCoV, using the S-glycoprotein as a model to shed light on the host-pathogen interaction in the dynamic process of SARS-CoV-2 infection. Although the amino acid sequences of the S-glycoprotein were different between the various HCoVs, the structural analysis exhibited high similarity; the best 3D structural overlap was found for SARS-CoV and SARS-CoV-2, consistent with the shared ACE2 predicted receptor.

Of note, the newly discovered SARS-CoV-2 genome has revealed differences between SARS-CoV-2 and SARS or SARS-like coronaviruses [28]. Although no amino acid substitutions were present in the receptor-binding motifs, that directly interact with human receptor ACE2 protein in SARS-CoV, six mutations occurred in the other region of the RBD [28, 29] were identified. On the other hand, the genomic comparative analysis highlighted the a strong diversity in the S gene among CoV in different hosts, confirming the biologically vital role of the S-glycoprotein as a key factor in viral entry in cross-species transmission events [30].

In addition, the comparative 3D structural data may facilitate the definition of already known antibody epitopes in the S-glycoprotein of other coronaviruses, it will also be useful in rational vaccine design and in gauging anti-virus directed immune responses after vaccination [27]. In fact, S-glycoprotein remains an important target for vaccines and drugs. previously evaluated in SARS and MERS, while a neutralizing antibody targeting the S-glycoprotein protein could provide passive immunity. The host interactome, linked to S-glycoprotein of SARS-CoV and MERS-CoV, mainly highlighted innate immunity pathway components, such as Toll Like receptors, cytokines and chemokines. The 3D structural analysis confirmed that we established that S-glycoprotein of SARS-CoV-2 has strong similarity in the 3D structure with SARS-CoV [16].

### Host interactome in all three HCoV infections

The reconstruction of virus-host interactome was carried out using RWR algorithm to explore the human PPI network and studying PPI and COEX multilayer. The PPI network topology of host interactome in all three infections indicated the presence of several hub proteins. In the HCoV-229E - host interactome hub position was hold by RAD18 and APEX, which play an important role in DNA repair due to UV damage in phase S [31].

For the SARS-CoV interactome, the gene hubs were identified in ACE2,CLEC4G and CD209, which are known interactors with S-glycoprotein of SARS-CoV[32, 33].

In fact, two independent mechanisms were described as trigger of SARS-CoV infection: proteolytic cleavage of ACE2 and cleavage of S-glycoprotein. The latter activates the glycoprotein for cathepsin L-independent host cell entry. Activated the S-glycoprotein by cathepsin L mechanism in host cell entry was reported in many infection of CoV, such as HCoV-229E and SARS-CoV [34, 35]. A recent study speculated that this interaction will be preserved in SARS-CoV-2 [17], but might be disrupted of a substantial number of mutations in the receptor binding site of S gene will occur. Likewise, the S-glycoprotein in SARS-CoV-2 is expected to interact with type II transmembrane protease (TMPRSS2) and probably is involved in inhibition of antibody-mediated neutralization [36, 37]. It is rather unexpected that, for this virus, no intracellular pathways were highlighted by the multilayer analysis, suggesting that this field is still open to further investigation.

In MERS-CoV infection a gene hub role was described for DPP4, which is known to regulate cytokine levels through catalytic cleavage [38]. Immune cell–recruiting chemokines and cytokines, such as IP-10/CXCL-10, MCP-1/CCL-2, MIP-1α/CCL-3, RANTES/CCL-5, can be strongly induced by MERS-CoV, showing higher inducibility in human monocyte–derived macrophages by MERS-CoV as compared to than SARS-CoV infection [39].

Finally, biological pathways, revealed by enrichment analysis in over all models, supported early activation of innate immune system, as Toll Like receptor Cascade and TGF-β for SARS-CoV, or chemokine and cytokine pathways and infection-related pathways for MERS-CoV, with a strong significance for both.

### Pathogenic model for HCoV infections

We constructed a host molecular interactome with SARS-CoV, MERS-CoV and HCoV-229E in patients with cancer, assuming that most of these interactions, especially for SARS-CoV, are shared with SARS-CoV-2. A network-based methodology, along with guilt-by-association algorithm (RWR), was applied to define the pathological model of COVID-19 and provide a treatment of SARS-CoV-2, using existing transcriptomic and proteomic information.

Based on the main pathways identified by the network-based interactome analysis, the following issues require focus further study:

#### First

The predicted receptor for SARS-CoV-2 has been inferred to be ACE-2, i.e. the same used by SARS-CoV, based on the high similarity of the S-glycoprotein of the two viruses, and this is the basis for hypothesizing to use SARS-CoV as a model for virus-host interactome in COVID-19;

#### Second

Mitogen activated protein kinase (MAPK) is a major cell signalling pathway that is known to be activated by diverse groups of viruses, and plays an important role in cellular response to viral infections. MAPK interacting kinase 1 (MNK1) has been shown to regulate both cap-dependent and internal ribosomal entry sites (IRES)-mediated mRNA translation;

#### Third

The identification of the MAPK pathway in SARS-CoV model is highly consistent with in vivo model, where P38 MAPK was found increased in the lungs of mice infected with SARS-CoV [40];

#### Fourth

The identification of the TGF-β pathway in S-glycoprotein-induced interactome for SARS-CoV of particular interest, due to the previous evidence that this virus, and in particular its protease, triggered the TGF-β through the p38 MAPK/STAT3 pathway in alveolar basal epithelial cells [41, 42];

#### Fifth

Innate immune pathways were identified in S-glycoprotein-induced models of SARS-CoV and MERS-CoV, as Toll Like receptor, cytokine and chemokine.

Every described pathway can be matched with clinical aspects, the data presented in this report may therefore aid to design a ‘blue print’ for SARS-CoV-2 associated pathogenicity.

The severity and the clinical picture of SARS-CoV and MERS-CoV infections could be related to the activation of exaggerated immune mechanism, causing uncontrolled inflammation [43]; however, the role of strong immune response in SARS-CoV-2 infection severity is still uncertain.

However, we may consider that host kinases link multiple signalling pathways in response to a broad array of stimuli, including viral infections. TGF-β, produced during the inflammatory phase by macrophages, is an important mediator of fibroblast activation and tissue repair. High levels of systemic inflammatory cytokines/chemokines has been widely reported for MERS-CoV infections [44–46], correlating with immunopathology and massive pulmonary infiltration into the lungs [47]. Also the HCoV-229E infection can be described with this distance model, although this infection was not associated with a severe respiratory disease. In fact, HCoV-229E is responsible for mild upper respiratory tract infections, such as common colds, with only occasional spreading to the lower respiratory tract, but it interacts with dendritic cells in the upper respiratory tract, inducing a cytopathic effect [48].

## Conclusions

In conclusion, we developed a network-based model, which could be the framework for structure-guided research process and for the pathogenetic evaluation of specific clinical outcome. Accurate structural 3D protein models and their interaction with host receptor proteins can allow to build a more detailed theoretical disease model for each HCoV infection, and support the drawing of a disease model for COVID-19. Our analyses suggests it is important to carry out *in silico* experiments and simulations through specific algorithms.

## Limitations to our study

A single protein, namely S-glycoprotein was used as seed, therefore the highlighted interactions were limited to those connected with this unique viral protein. However, this is a proof of concept study, from which it appears that a similar approach may be used to study other viral proteins interacting with host cell pathways. Another limitation is that the pathway analysis did not consider tissue and cell type diversity. Finally, the low threshold established for the number of nodes found by RWR (200) limited the reconstruction of the entire pathways. However, this was a software-imposed threshold.

In summary, the interactome analysis aided to guide the design of novel models of SARS-CoV pathogenicity.

## Abbreviations

CoVs: Coronaviruses
HCoV: Human Coronavirus
PPI: Protein-Protein Interactions
COEX: Gene Coexpression data
SARS-CoV: Severe Acute Respiratory Syndrome Coronavirus
MERS-CoV: Middle East Respiratory Syndrome Coronavirus
S-glycoprotein: Spike glycoprotein
RWR: Random walk with restart
FDR: False Discovery Rate

## Authors’ contributions

Conceptualization: F.M., E.G., F.N.L. Data curation: F.M., E.G., Formal analysis: F.M., E.G., Funding acquisition: M.R.C., G.I., F.V. Investigation: F.M., E.G., F.N.L. Methodology: F.M., E.G., T.A., S.A. Resources: M.R.C., G.I. Software: F.M., E.G. Supervision: M.R.C., F.N.L. Validation: C.A., F.V., G.K., M.M., M.P. Visualization: F.M., E.G. Writing ± original draft: F.M., E.G., M.R.C., F.N.L., G.K., A.Z., M.M. Writing ± review & editing: F.M., E.G., M.R.C., F.N.L. A.Z. All authors reviewed and approved the manuscript.

## Acknowledgments

We gratefully acknowledge: Collaborators Members of INMI COVID-19 study group; COVID 19 INMI Network Medicine for IDs Study Group:

Abbate Isabella, Agrati Chiara, Al Moghazi Samir, Ascoli Bartoli Tommaso, Capobianchi Maria Rosaria, Capone Alessandro, Goletti Delia, Rozera Gabriella, Nisii Carla, Gagliardini Roberta, Ciccosanti Fabiola, Fimia Gian Maria, Nicastri Emanuele, Giombini Emanuela, Lanini Simone, D’Abramo Alessandra, Rinonapoli Gabriele, Girardi Enrico, Bartolini Barbara, Montaldo Chiara, Marconi Raffaella, Antonio Addis, Bradley Maron, Bianconi Ginestra, De Meulder Bertrand, Kennedy Jason, Khader Shabaana Abdul, Luca Francesca, Maeurer Markus, Piacentini Mauro, Merler Stefano, Pantaleo Giuseppe, Rafick-Pierre Sekaly, Sanna Serena, Segata Nicola, Zumla Alimuddin, Messina Francesco, Lauria Francesco Nicola, Ippolito Giuseppe, Vairo Francesco.

## Consent for publication

Not applicable.

## Ethics approval and consent to participate

Not applicable.

## Availability of data and materials

PPI data of SARS-CoV, MERS-CoV, HCoV-229E S-glycoprotein were inferred through published PPI data, using STRING Viruses (http://viruses.string-db.org/) and VirHostNet (http://virhostnet.prabi.fr/), as well as published scientific reports with a focus on virus-host interactions [18–20]. Human PPI databases (BioGrid, InnateDB-All, IMEx, IntAct, MatrixDB, MBInfo, MINT, Reactome, Reactome-FIs, UniProt, VirHostNet, BioData, CCSB Interactome Database), using R packages PSICQUIC (https://bioconductor.org/packages/release/bioc/html/PSICQUIC.html) and biomaRt (https://bioconductor.org/packages/release/bioc/html/biomaRt.html) [21, 22]. The gene expression data set was built from the Protein Atlas database (https://www.proteinatlas.org/) [23].

---

Figure S1. Pairwise distances along 259 full length CoV genomes. In the bottom of picture, indicative gene positioning along CoVs genomes is reported. The list of all considered genomes is reported in Table S1.

Figure S2. 3D structure of S-glycoprotein of SARS-CoV-2 and comparison with the ortholog from HCoV-229E, SARS-CoV, and MERS-CoV. Lateral (a) and superior (b) representation of SARS-CoV-2 S-glycoprotein, deducted for the sequence of patient INMI1 (MT066156.1). Each subunit chain has a different color. Structure comparison of S-glycoprotein subunit between: HCoV-229E and SARS-CoV-2, in purple and blue respectively (c); SARS-CoV and SARS-CoV-2, in red and blue, respectively (d); MERS-CoV and SARS-CoV-2, in green and blue, respectively (e).

Figure S3.Amino acid alignment and secondary motifs in the receptor binding domain (RBD) of S-glycoprotein of HCoV-229E, SARS-CoV, MERS-CoV and SARS-CoV-2 are shown. Legend of secondary motifs identifiers: H = α Helix, E = β Sheet, X = Random coil.

Figure S4. HCoV-229E - host interactome resulting from RWR applied to the top 200 closest proteins identified by RWR, using S-glycoprotein of HCoV-229E.

Figure S5. SARS-CoV - host interactome resulting from RWR applied to the top 200 closest proteins identified by RWR, using S-glycoprotein of SARS-CoV.

Figure S6. MERS-CoV - host interactome resulting from RWR applied to the top 200 closest proteins identified by RWR, using S-glycoprotein of MERS-CoV.

Table S1: List of accession numbers of H-CoV.

Table S2: list of genes selected by RWR algorithm for HCoV-229E, along with proximity score.

Table S3: list of genes selected by RWR algorithm for SARS-CoV, along with proximity score.

Table S4: list of genes selected by RWR algorithm for MERS-CoV, along with proximity score.

